# Pharmacologic METTL3 inhibition attenuates TGF-β–driven profibrotic signature in systemic sclerosis

**DOI:** 10.64898/2026.09.10.750422

**Authors:** Nikolaos I. Vlachogiannis, Simon Tual-Chalot, Maria Polycarpou-Schwarz, Elly Wu, Aikaterini-Paraskevi Avdi, Kateryna Sopova, Marco Sachse, Stylianos Panopoulos, Maria Tektonidou, Kimon Stamatelopoulos, Athanasios Zovoilis, Petros P. Sfikakis, Konstantinos Stellos

## Abstract

**One Sentence Summary:** Patients with systemic sclerosis carry a distinct m6A RNA methylation profibrotic signature, that can be reversed by METTL3 inhibitors.

Systemic sclerosis (SSc) is a devastating fibrosing disorder affecting the skin and internal organs, associated with the highest mortality among all rheumatic diseases. The mechanisms underlying excessive fibrogenesis in SSc are incompletely defined and treatment options are limited. N6-methyladenosine (m6A), the most abundant internal mRNA modification, is a reversible and pharmacologically tractable determinant of RNA fate. Here we mapped the m6A epitranscriptome of peripheral blood mononuclear cells (PBMCs) from patients with SSc using two orthogonal single-nucleotide-resolution platforms, a MazF-based m6A microarray and nanopore direct RNA sequencing. SSc PBMCs showed coordinated upregulation of the m6A writer complex (METTL3, METTL14, WTAP), a two-fold increase in global m6A, and redistribution of m6A marks from 3′ untranslated regions toward coding sequences. Both platforms converged on TGF-β signaling as the most enriched pathway among differentially methylated transcripts, with hypermethylation of *TGFB1*, *JUNB* and *KLF10* transcripts. Pharmacologic inhibition of METTL3 with STM2457 significantly reversed TGF-β-induced transcriptomic changes in skin fibroblasts preventing their transformation into pathological myofibroblasts. Together, these findings uncover an epitranscriptomic signature driving profibrotic transcriptional alterations in SSc, including enhanced TGF-β signaling, which can be effectively reversed by clinically available enzymatic m6A inhibitors. Our data suggest that targeting m6A methylation may offer a novel therapeutic strategy to mitigate fibrosis in SSc.

## INTRODUCTION

Systemic sclerosis (SSc) is a prototypical fibrosing disease affecting the skin and internal organs, characterized by the unique pathogenetic triad of microvasculopathy, (auto)immune activation and fibrosis [1]. Persistent activation of fibroblasts and their differentiation into extracellular matrix (ECM)– producing myofibroblasts represent central pathogenic events driving tissue remodeling and organ fibrosis in SSc [2]. Transforming growth factor–β (TGF-β) signaling is a core regulator of this process and strongly correlates with disease severity and progression [3–5], making this pathway a focal point for mechanistic and therapeutic studies [6–8]. Nonetheless, the upstream mechanisms that sustain persistent TGF-β pathway activation in SSc remain poorly defined.

N6-methyladenosine (m6A) is the most abundant internal modification in human mRNA [9–11], dynamically shaping the transcriptome and influencing RNA fate in a context-dependent manner [10]. m6A is catalyzed by a multicomponent ‘writer’ complex composed of the methyltransferases METTL3, the main catalytic enzyme, and METTL14, along with essential adaptor proteins such as WTAP [12] [13]. Dynamic demethylation is carried out by the ‘eraser’ enzymes FTO and ALKBH5, conferring reversibility to m6A marks and making this pathway pharmacologically tractable. Through the recruitment of specific ‘reader’ proteins, m6A regulates nearly every aspect of RNA metabolism, including splicing [14], nuclear export, stability [15,16], processing [17], and translation [18–21]. Recent advances in m6A detection technologies, such as direct long-read RNA sequencing [22] and antibody-based enrichment methods (MeRIP-seq, m6A-seq) [23,24], have uncovered critical roles for m6A in diverse disease contexts, including chronic inflammatory and fibrotic disorders [9]. In particular, m6A has been shown to modulate fibrotic signaling pathways in the liver [25], kidneys [26] and lungs [27], positioning it as a potential therapeutic target in fibrosis.

Emerging evidence supports a strong interplay between m6A methylation and TGF-β signaling at multiple levels. m6A regulates *TGFΒ1* mRNA expression [28], as well as transcript stability and translation of downstream effectors such as *JUN* and *JUNB* [29] [30]. On the other hand, TGF-β regulates the m6A methylome via SMAD2/3-mediated recruitment of the m6A writer complex to distinct transcripts [31]. We therefore hypothesized that epitranscriptomic dysregulation of the m6A methylome contributes to persistent TGF-β activation and fibrotic gene expression in SSc. We provide the first transcriptome-wide map of the m6A methylome in peripheral blood mononuclear cells (PBMCs) from patients with SSc and examine the effects of pharmacologic m6A inhibition on TGF-β– driven profibrotic gene expression in skin fibroblasts. Together, our findings identify m6A signaling as a previously unrecognized and therapeutically targetable regulator of fibrotic responses in SSc.

## RESULTS

### Increased m6A ‘writers’ expression and global m6A methylation levels in SSc PBMCs

To determine whether the m6A epitranscriptomic machinery is dysregulated in SSc, we first quantified the expression of core m6A writer complex components and global m6A methylation levels in PBMCs from SSc patients and healthy controls (demographics of the cohort have been published in our previous work [32]). Expression of the methyltransferases *METTL3* and *METTL14*, and the auxiliary protein WTAP was increased by approximately 1.5-fold in SSc PBMCs (n=31) compared to healthy individuals (n=24) (all P<0.001; **Fig. 1A-D**). Measurement of global m6A levels in a sub-cohort with available RNA consisting of 9 SSc patients *vs* 9 HC revealed a 2-fold increase of m6A in SSc PBMCs (**Fig. 1E**), which correlated with *METTL3* (n=18, r=0.446, P=0.06; **Fig. 1F**), *METTL14 (*n=18, r=0.486, P=0.04; **Fig. 1G**) and *WTAP* expression (n=18, r=0.550, P<0.001; **Fig. 1H**).

**Fig. 1.**
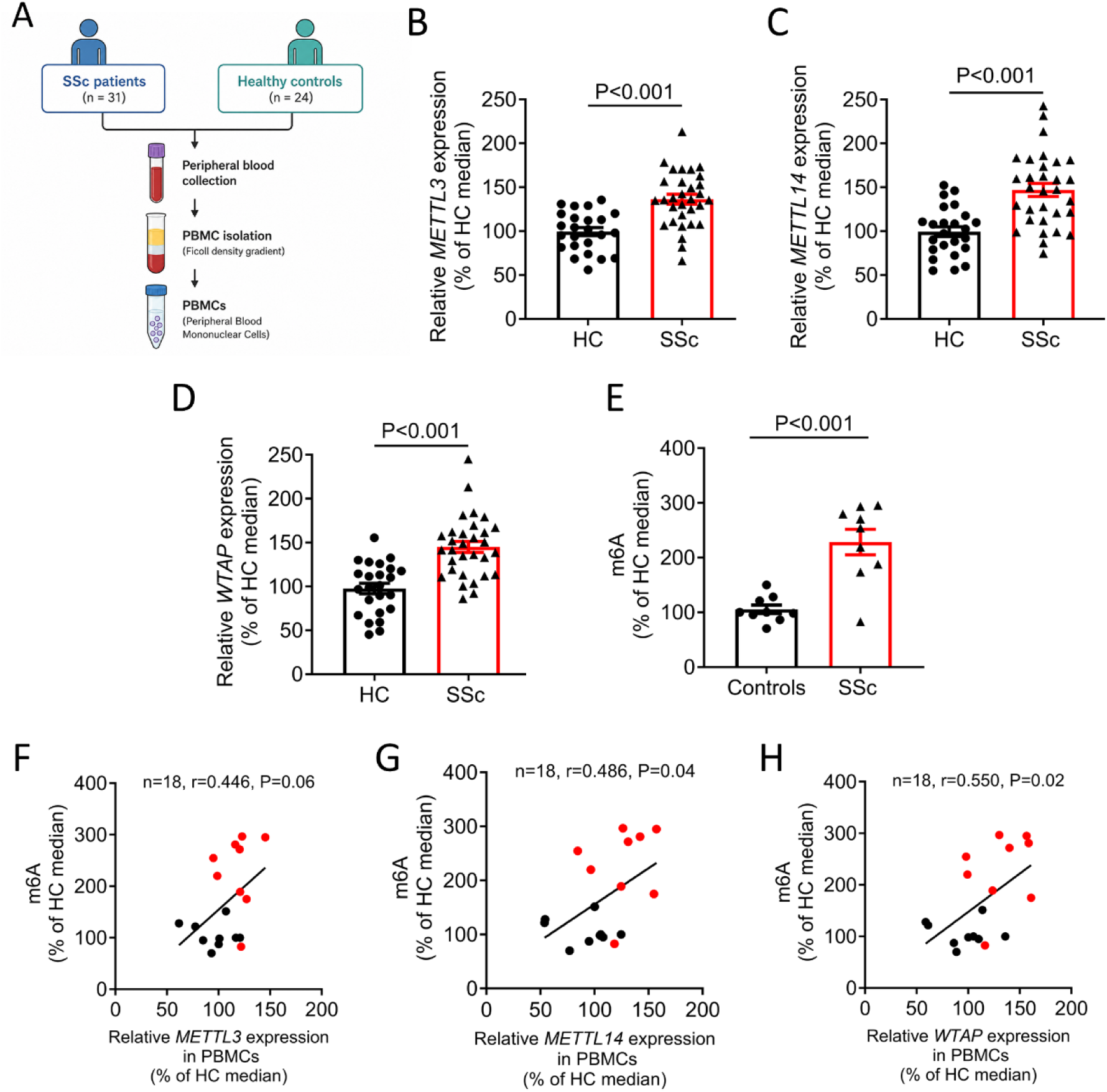
Expression of m6A writer complex components and global m6A levels in SSc PBMCs. **A-D.** Relative mRNA expression of the components of the m6A writer complex, namely *METLL3* (A), *METTL14* (B) and the adaptor protein *WTAP* (C), as quantified using qRT-PCR in PBMCs derived from 31 SSc patients and 24 apparently healthy controls (HC). **E.** Total m6A methylation levels in PBMCs of 9 SSc patients *vs* 9 healthy controls as quantified by EpiQuik ™ m6A RNA Methylation Quantification Kit (P-9005, EpiGenTek). **F-H**. Scatterplots showing the correlation of m6A methylation levels with expression *METTL3* (E), *METTL14* (F)*, WTAP* (G). Bar-graphs (A-D) represent mean ± standard error of the mean. P-values are derived from independent samples t-test or Welch’s t-test, where applicable. Correlation co-efficient (E-G) was determined using the Pearson’s test. Bar-graphs represent mean ± standard error of the mean.

### Transcriptome-wide site-specific m6A methylation analysis

To map the m6A methylome at single-nucleotide resolution, we performed transcriptome-wide m6A profiling of PBMCs from 10 SSc patients and 3 HC using the Arraystar m6A Single Nucleotide Microarray platform, which enables simultaneous quantification of methylation stoichiometry across thousands of known m6A sites (**Fig. 2A**, **Table 1**). Of the 11,237 single methylation sites included in the array, 10,123 unique m6A sites mapping on 4,726 unique genes were detected in our samples; 9,895 (97.75%) m6A sites were mapped on protein coding transcripts, while the remaining 228 (2.25%) m6A sites were mapped on non-coding transcripts (**Fig. 2B**). Among the m6A sites on coding transcripts, 305 (3.08%) were detected in 5’ untranslated region (UTR), 5,784 (58.45%) in coding regions, 3,806 (38.46%) in 3’ UTR and the rest mapped in other gene regions (**Fig. 2B**). Principal component analysis (PCA) of m6A abundance in the 10,123 unique m6A sites revealed a clear demarcation between SSc patients and HC (**Fig. 2C**). Applying a threshold of 1.5-fold absolute change in m6A methylation and P<0.05 we were able to identify 491 hyper-methylated sites mapping on 460 unique genes, and 639 hypo-methylated sites mapping on 568 unique genes (**Fig. 2C, D, Supplementary Table 1**). Pathway enrichment analysis of differentially methylated genes using Ingenuity Pathway Analysis revealed TGF-β signaling as the most enriched pathway **(Figure 2E**), consistent with a central role of aberrant m6A deposition in sustaining profibrotic signalling in SSc. Differentially methylated sites between SSc and HC mapping in genes belonging to the TGF-β pathway are shown in **Fig. 2F**.

**Fig. 2.**
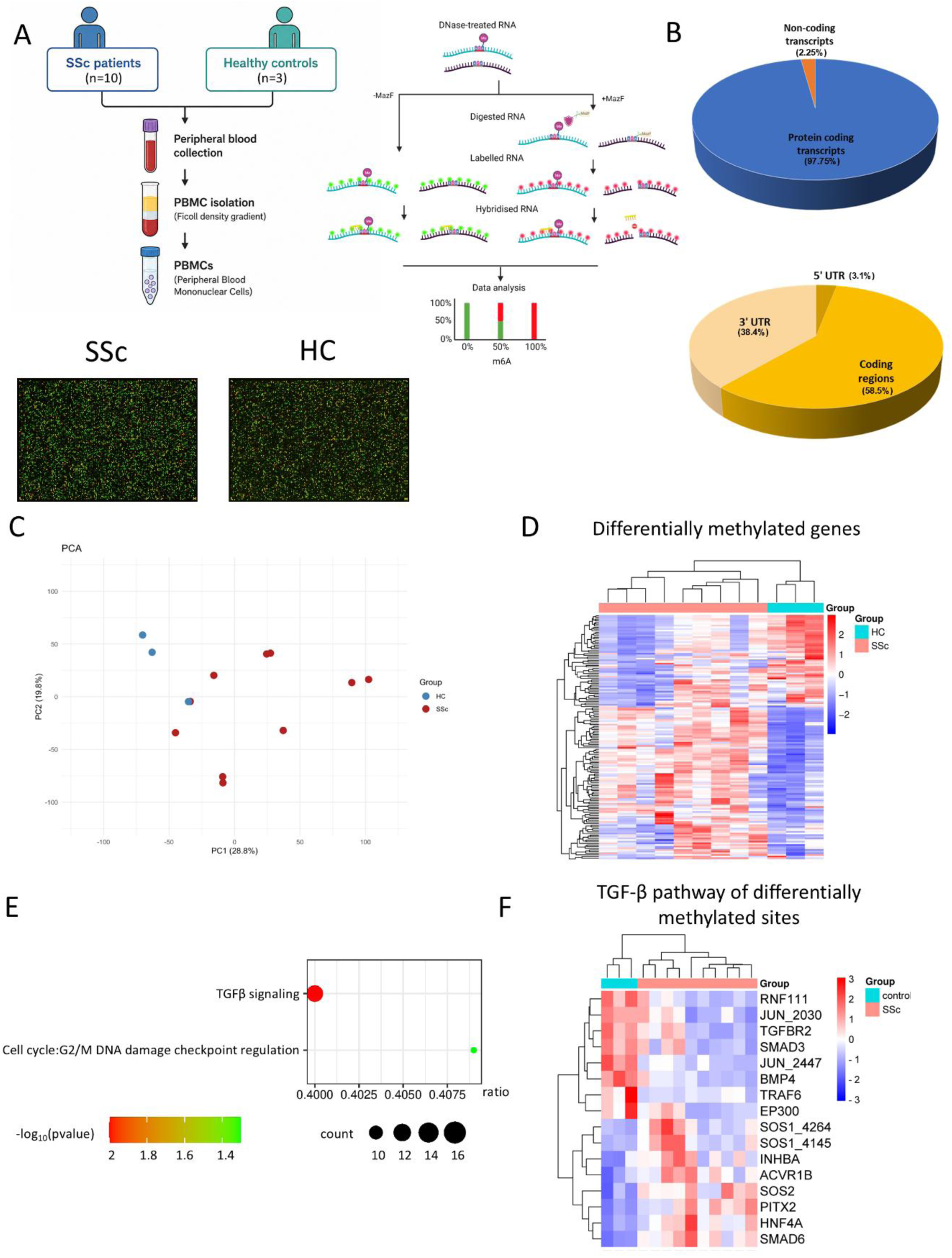
Differential methylome analysis in SSc vs HC PBMCs using an array-based single-base m6A quantification assay. **A.** Study design of the methylome study using Arraystar m6A Single Nucleotide Microarray. **A.** Pie chart of the distribution of the methylated site identified. **C.** Principal component analysis of the m6A methylome in 10 SSc patients and 3 age- and sex-matched healthy individuals. Absolute methylation quantification in 10,123 unique m6A sites was used as input. **D.** Heatmap of differentially methylated genes in SSc vs HC PBMCs (491 hyper-methylated sites mapping on 460 unique genes, and 639 hypo-methylated sites mapping on 568 unique genes). A cut-off of 1.5-fold absolute fold-change in m6A methylation rate and P<0.05 was used. **E.** Ingenuity pathway analysis of differentially methylated genes in SSc vs HC PBMCs. **F.** Heatmap of differentially methylated sites belonging to components of the TGF-β pathway in SSc vs HC PBMCs.

**Table 1.**
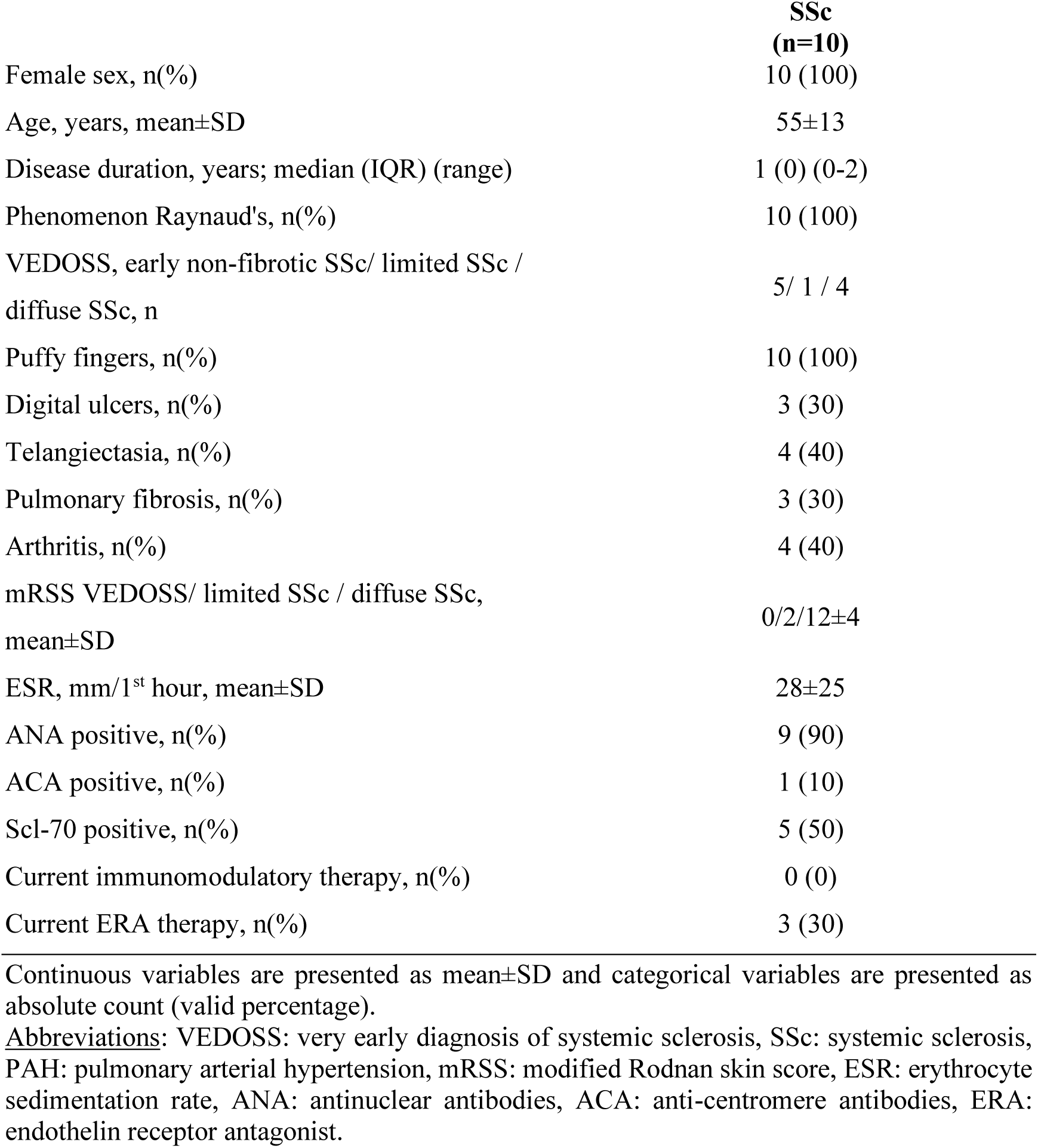
Demographics and disease characteristics of the SSc Arraystar cohort.

|  | **SSc**<br>**(n=10)** |
| --- | --- |
| Female sex, n(%) | 10 (100) |
| Age, years, mean±SD | 55±13 |
| Disease duration, years; median (IQR) (range) | 1 (0) (0-2) |
| Phenomenon Raynaud's, n(%) | 10 (100) |
| VEDOSS, early non-fibrotic SSc/ limited SSc /<br>diffuse SSc, n | 5/ 1 / 4 |
| Puffy fingers, n(%) | 10 (100) |
| Digital ulcers, n(%) | 3 (30) |
| Telangiectasia, n(%) | 4 (40) |
| Pulmonary fibrosis, n(%) | 3 (30) |
| Arthritis, n(%) | 4 (40) |
| mRSS VEDOSS/ limited SSc / diffuse SSc,<br>mean±SD | 0/2/12±4 |
| ESR, mm/1<sup>st</sup> hour, mean±SD | 28±25 |
| ANA positive, n(%) | 9 (90) |
| ACA positive, n(%) | 1 (10) |
| Scl-70 positive, n(%) | 5 (50) |
| Current immunomodulatory therapy, n(%) | 0 (0) |
| Current ERA therapy, n(%) | 3 (30) |

Continuous variables are presented as mean±SD and categorical variables are presented as absolute count (valid percentage).
Abbreviations: VEDOSS: very early diagnosis of systemic sclerosis, SSc: systemic sclerosis, PAH: pulmonary arterial hypertension, mRSS: modified Rodnan skin score, ESR: erythrocyte sedimentation rate, ANA: antinuclear antibodies, ACA: anti-centromere antibodies, ERA: endothelin receptor antagonist.

### Mapping the m6A methylome in SSc PBMCs using Nanopore direct RNA sequencing

Having established a disease-associated m6A signature by array-based profiling, we next sought to map m6A modifications at transcriptome-wide scale using an antibody-independent exploratory approach. We performed Oxford Nanopore direct RNA sequencing on PBMCs from SSc patients and healthy controls (HC) (n=3 per group), enabling direct detection of RNA modifications at single-nucleotide resolution without chemical conversion or immunoprecipitation (**Fig. 3A**). m6A methylation was predicted using m6Anet. Of 123,258 DRACH sites tested for m6A modification, 750 sites achieved a modification probability of >0.9. After further filtering for sites detected in more than two samples in at least one group and confirmed transcript expression, 645 high-confidence m6A sites were retained for downstream analysis (**Supplementary Table 2**). Principal component analysis and unsupervised hierarchical clustering revealed a distinct m6A methylation profile separating SSc patients from HC (**Fig. 3B, C**), corroborating the disease-associated epitranscriptomic signature observed with the Arraystar platform. In HC, high-confidence m6A sites followed the canonical distribution, with enrichment around stop codons and in 3′ untranslated regions (3′UTRs), regions classically associated with post-transcriptional regulation of transcript stability [23,33]. In contrast, SSc samples displayed a marked redistribution of m6A marks, with increased methylation in coding regions at the expense of 3′UTRs (**Fig. 3D, E**), suggesting a disease-specific reprogramming of the epitranscriptomic landscape. Motif analysis of high-confidence m6A sites identified GGACT as the predominant DRACH motif in both groups, with no significant differences in motif composition between HC and SSc (**Fig. 3F**).

**Fig. 3.**
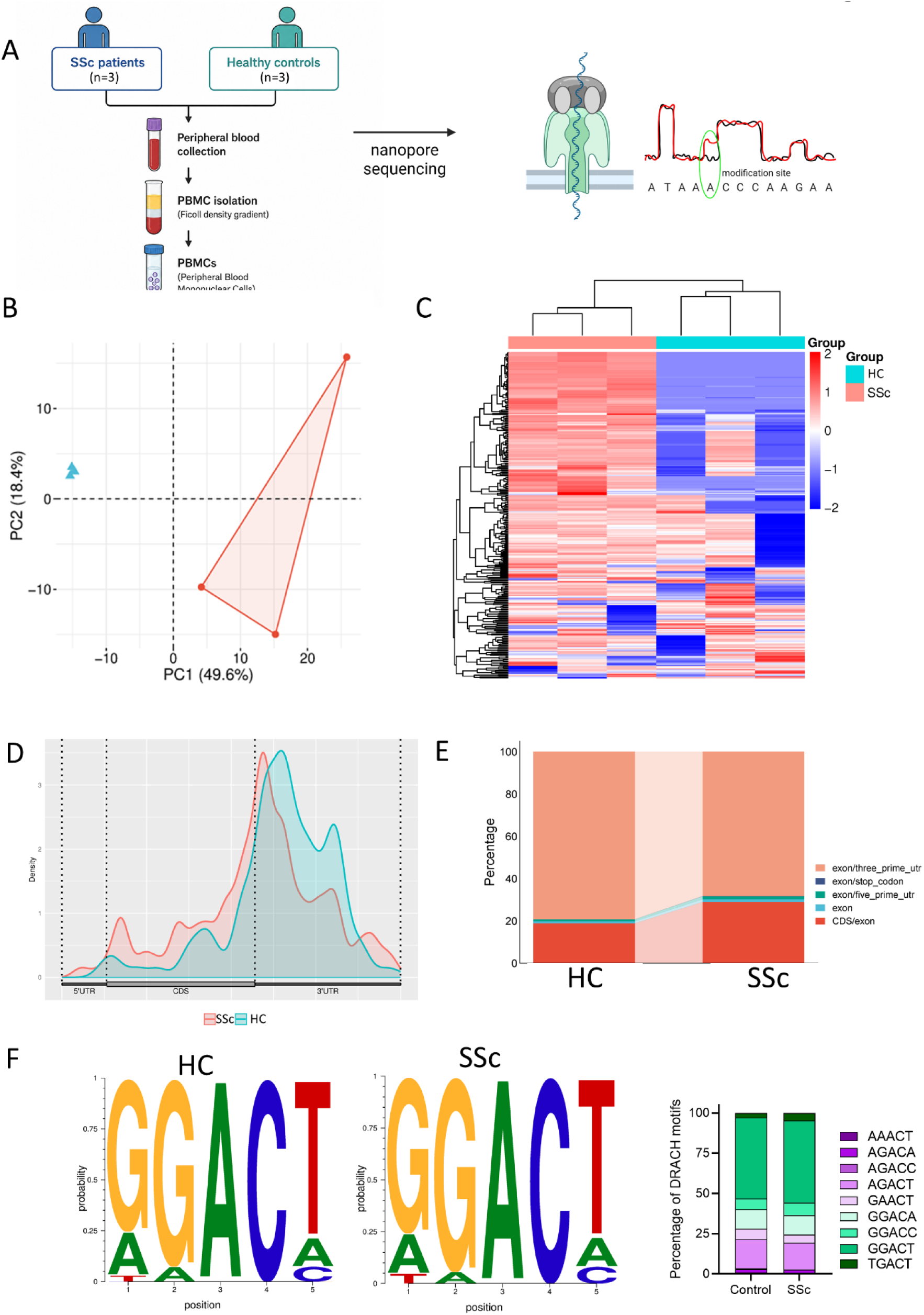
Mapping of the m6A methylome with Nanopore direct RNA-seq. **A.** Experimental workflow for Nanopore direct RNA sequencing of RNA from PBMCs of 3 systemic sclerosis (SSc) patients and 3 age- and sex-matched healthy controls (HC), including signal-level alignment and m6A calling with m6Anet to define high-confidence m6A sites. **B.** Principal component analysis (PCA) of m6A methylation profiles across 645 high-confidence m6A sites identified. **C.** Heatmap of high-confidence methylated sites validated in SSc vs HC PBMCs. **D.** Metagene plot showing the distribution of m6A modification sites along mRNA isoforms for HC and SSC individuals. **E.** Distribution of m6A sites across 5′ UTRs, CDS, and 3′ UTRs. **F.** Motif analysis of validated m6A sites and quantification of the proportion of each DRACH motif observed in high-confidence m6A sites.

### Differential m6A methylome analysis in SSc PBMCs reveals enrichment of profibrotic pathways

Having characterized the global m6A landscape, we next performed differential methylation analysis to identify site-specific m6A changes between SSc patients and HC. A total of 502 hypermethylated and 107 hypomethylated sites were detected across the transcriptome, corresponding to 212 hypermethylated and 33 hypomethylated genes at the gene level (**Fig. 4A**). The predominance of hypermethylated over hypomethylated sites was consistent with the upregulated expression of multiple genes encoding m6A writers and readers observed in SSc PBMCs (**Fig. 4B**), while the increase in m6A methylation moderately correlated with increased expression of the corresponding transcript (r=0.33, P<0.0001; **Fig. 4C**).

**Fig. 4.**
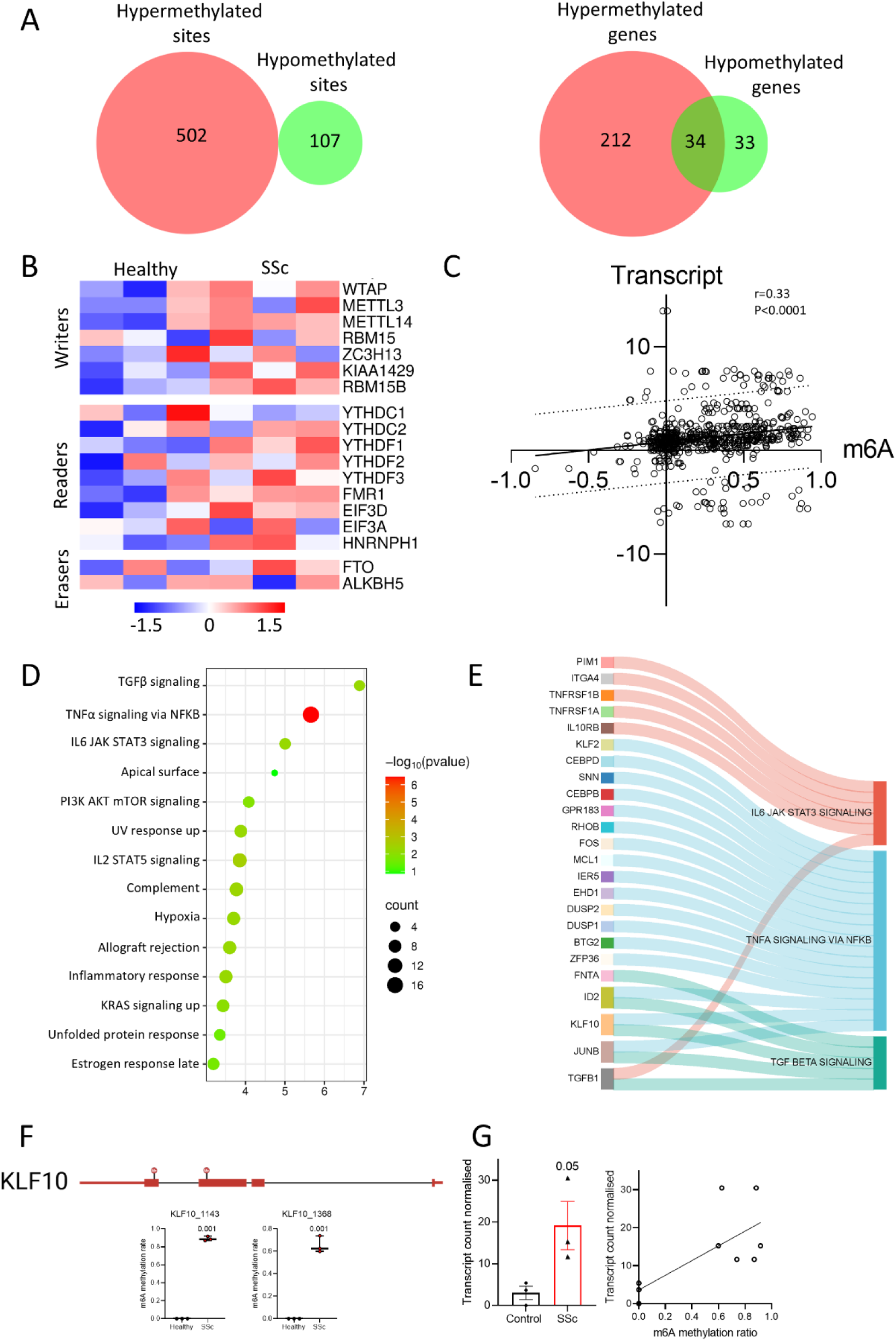
Enrichment analysis of differentially methylated sites in SSc vs HC PBMCs using Nanopore. **A.** Venn diagrams showing intersections of hypermethylated and hypomethylated high-confidence m6A sites and their genes. **B**. Relative expression of selected genes encoding components of the m6A machinery (writers, readers, and erasers) in SSc versus HC PBMCs. **C.** Correlation between changes in m6A methylation and transcript abundance for genes harboring at least one differentially methylated site. **D.** Pathway enrichment analysis of genes with exclusively hypermethylated m6A sites. **E.** Sankey dot plot visualising the genes from the top 3 enriched pathway. **F-G.** m6A modification along the KLF10 transcript in HC and SSc PBMCs, illustrating that an m6A site within the coding region is detected exclusively in SSc samples and is associated with increased KLF10 transcript abundance in SSc.

To identify dysregulated biological pathways, we performed Ingenuity Pathway Analysis on the 212 genes exclusively hypermethylated sites. The three most significantly enriched pathways were “TGF-β signaling”, “TNFα signaling via NF-κB”, and “IL-6/JAK/STAT3” signaling (**Fig. 4D**), findings that were concordant with the pathway enrichment results obtained from the Arraystar platform, providing cross-platform validation of the epitranscriptomic signature. Among the differentially methylated genes within the TGF-β signaling pathway, *FNTA*, *ID2*, *KLF10*, *JUNB*, and *TGFB1* were identified as hypermethylated in SSc PBMCs (**Fig. 4E**). Mapping of individual m6A sites across these transcripts revealed that KLF10 methylation was detected exclusively within the coding region of SSc samples and was absent in HC (**Fig. 4F**). This coding-region hypermethylation was associated with increased KLF10 transcript abundance in SSc patients (**Fig. 4G**), suggesting that m6A deposition may enhance KLF10 expression through improved transcript stability or translational efficiency. Notably, KLF10 has recently been identified as a member of an inflammatory gene module, together with JUNB, in monocytes from the peripheral blood of SSc patients [34], supporting its relevance to SSc immunopathology. *FNTA*, *ID2*, *JUNB*, and *TGFB1* also showed an overall trend toward hypermethylation across their detected m6A sites (**Supplementary Fig. 1**).

### METTL3 inhibition attenuates TGF-β-induced transcriptomic reprogramming and profibrotic gene expression in fibroblasts

To examine the functional consequences of m6A dysregulation on TGF-β-induced profibrotic responses, we co-treated healthy human dermal fibroblasts with TGF-β1 and STM2457, a selective enzymatic inhibitor of METTL3, a derivative of which is currently being evaluated in clinical trials [35] (**Fig. 5A**).

**Fig. 5.**
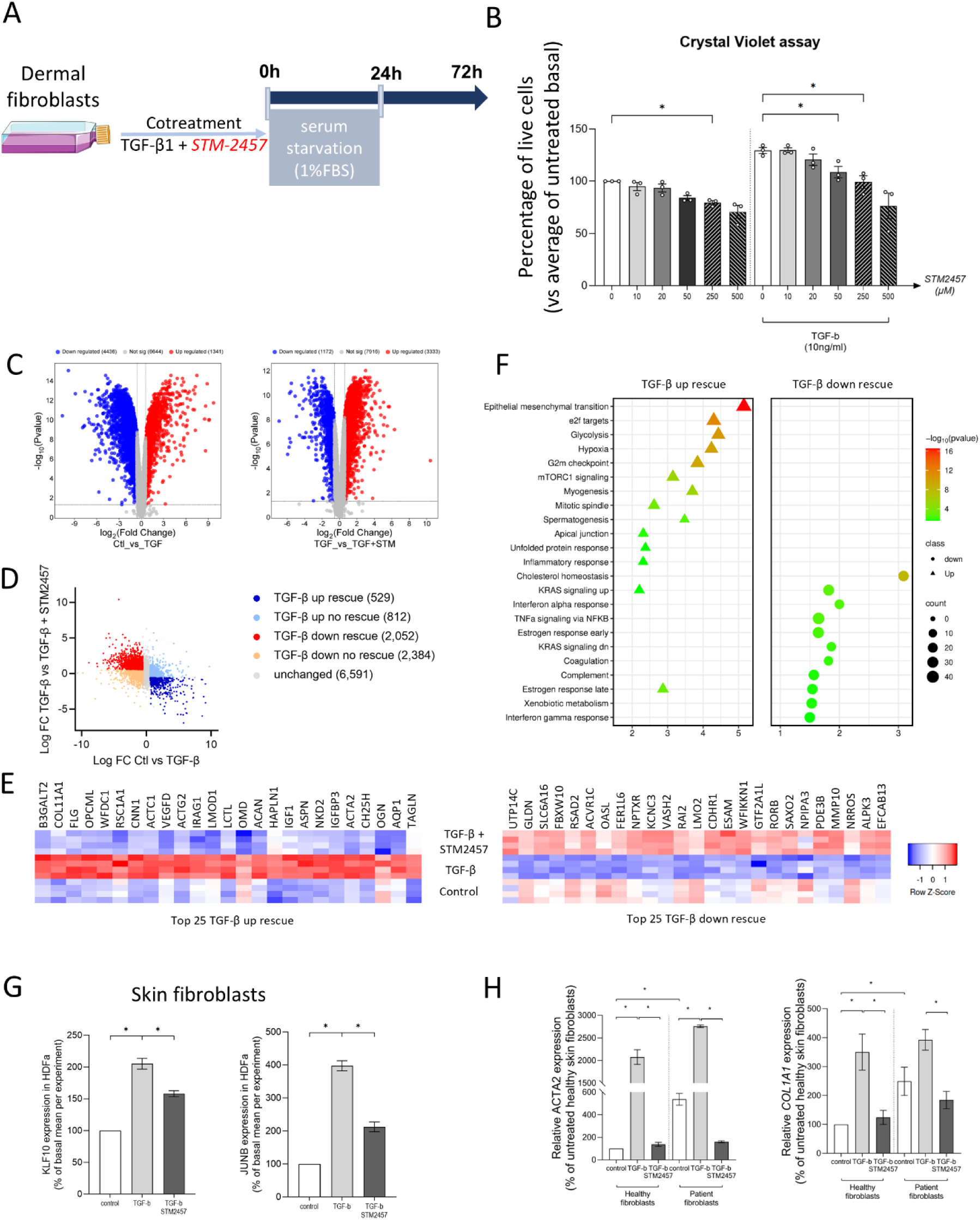
Enzymatic inhibition of METTL3 ameliorates TGF-β-induced gene expression in dermal fibroblasts. **A.** Study design of the *in vitro* treatment of healthy skin fibroblasts (HDFa) with TGF-β1 and the enzymatic inhibitor of METTL3, STM2457. **B.** Viability of skin fibroblasts treated with increasing concentrations of STM2457. Bar graphs mean ± SEM of 3 independent biological replicates, expressed as percentage of viable cells compared to untreated cells. **C**. Volcano plots displaying log₂(Fold Change) on the x-axis against −log₁₀(p-value) on the y-axis of the control versus TGF-β-treated cells (Ctl_vs_TGF) and treatment with STM in the context of TGF-β stimulation (TGF_vs_TGF+STM). **D**. Scatter plot identifying the STM2457 rescue signature. For a gene to be called “STM rescued”, it had to satisfy all three conditions: log₂(Fold Change) TGFβ vs Ctl > 0.58 and FDR < 0.05 (TGFβ changed it), log₂(Fold Change) TGFβ vs TGFβ+STM > 0.58 and FDR < 0.05 (STM reversed it), and the STM2457 log₂(Fold Change) direction must oppose the TGFβ direction. **E**. Heatmap of the top 25 genes rescue by STM2457 after upregulation or downregulation of TGF-β1. **F**. GSEA against the Hallmark gene set collection of the genes rescued by STM2457 after upregulation or downregulation og TGF-β1. **G.** Expression of *KLF10* and *JUNB* in healthy skin fibroblasts treated with TGF-β1 and STM2457 for 48h. **H.** Expression of *ACTA2* and *COL1A1* in healthy and patient-derived skin fibroblasts treated with TGF-β1 and STM2457 for 48h. Bar graphs mean ± SEM of 5 independent biological replicates.

Cell viability assessed by Crystal Violet assay confirmed that STM2457 was well tolerated across a wide concentration range (10–500 µM), both in the presence and absence of TGF-β1, with no significant cytotoxicity observed at the doses used for subsequent experiments (**Fig. 5B**).

To characterise the transcriptome-wide impact of METTL3 inhibition on TGF-β-driven gene expression, we performed RNA sequencing on healthy human skin fibroblasts treated with TGF-β1 alone, TGF-β1 & STM2457 or untreated (vehicle, control). Differential expression analysis identified 5,777 differentially expressed genes (DEGs) in the Ctl_vs_TGF-β comparison (4,436 downregulated, 1,341 upregulated), and 4,505 DEGs in the TGFb_vs_TGFb+STM2457 comparison (1,172 downregulated, 3,333 upregulated) (**Fig. 5C**).

To identify genes whose TGF-β1-induced expression changes were specifically reversed by METTL3 inhibition, we applied a stringent rescue criterion requiring that a gene: (i) was significantly altered by TGF-β1 relative to control (|log₂FC| >0.58, FDR <0.05); (ii) was significantly reversed by TGF-β1 + STM2457 relative to TGF-β1 alone (|log₂FC| >0.58, FDR <0.05); and (iii) the direction of the STM2457 effect opposed that of TGF-β1, confirming true rescue rather than additive dysregulation. Applying these criteria, we identified 529 TGF-β1-upregulated genes rescued by STM2457 (TGF-β up rescue) and 2,052 TGF-β1-downregulated genes rescued by STM2457 (TGF-β down rescue) (**Fig. 5D**). Among the top 25 TGF-β up rescue genes were established mediators of myofibroblast activation and ECM remodelling including *ACTA2*, *TAGLN*, *CNN1*, *ACTG2*, and *COL11A1* (**Fig. 5E**). Conversely, the top TGF-β down rescue genes included *RSAD2*, *OASL*, *MMP10*, and *RORB* (**Fig. 5E**), suggesting that METTL3 inhibition also partially restores innate immune and tissue homeostasis gene programmes suppressed by TGF-β1.

Among genes upregulated by TGF-β1 and rescued (downregulated) by STM2457 (TGF-β up rescue; **Fig. 5D**, left panel), the most significantly enriched pathway was “Epithelial-Mesenchymal Transition”, followed by “E2F targets”, “Glycolysis”, “Hypoxia”, and “G2M checkpoint. Among genes downregulated by TGF-β1 and rescued (upregulated) by STM2457 (TGF-β down rescue; **Fig. 5D**, right panel), the enriched pathways comprised “Cholesterol homeostasis”, “Interferon alpha response”, “TNFα signaling via NF-κB”, “Estrogen response early”, and “KRAS signaling”. Together, these findings indicate that STM2457 preferentially reverses the TGF-β1-driven mesenchymal, proliferative, and metabolic transcriptional programme, while restoring suppressed immune signalling gene programmes.

Consistent with these transcriptome-wide findings, targeted validation confirmed that METTL3 inhibition attenuated TGF-β1-induced expression of KLF10 and JUNB in cultured skin fibroblasts (**Fig. 5G**), two genes identified as hypermethylated in SSc PBMCs from the Nanopore sequencing analysis, directly linking the epitranscriptomic signature observed in patient samples to functional profibrotic responses in fibroblasts. Furthermore, METTL3 inhibition significantly attenuated TGF-β1-induced expression of ACTA2 and COL1A1, canonical markers of myofibroblast transformation, in both healthy and SSc-derived dermal fibroblasts (**Fig. 5H**), demonstrating that the antifibrotic effect of METTL3 inhibition extends to disease-relevant cellular contexts.

## DISCUSSION

In the current manuscript we provide the first comprehensive map of the m6A methylome in PBMCs of patients with SSc, the prototypic systemic fibrotic disorder. PBMCs from patients with SSc display increased expression of the core m6A writer complex components METTL3, METTL14 and WTAP, accompanied by elevated global m6A levels compared to healthy individuals. Using complementary approaches, we map transcriptome-wide m6A sites and identify differentially methylated transcripts within transcripts related to TGF-β signaling and fibroblast activation, supporting a central role of m6A in shaping the SSc profibrotic transcriptome. In dermal fibroblasts, catalytic inhibition of METTL3 with STM2457 reversed a defined subset of the TGF-β1–induced transcriptome and lowered TGF-β1– driven KLF10, JUNB, ACTA2 and COL1A1 in both healthy and patient-derived cells. Together these data position m6A as a layer of post-transcriptional control that reinforces TGF-β signaling in SSc.

Despite the growing recognition of m6A methylation role in various fibrotic disorders, its involvement in SSc has been largely unexplored until now. Using publicly available RNA-seq data from SSc skin samples, decreased expression of m6A erasers FTO and ALKBH5, as well as the adaptor protein WTAP, in SSc skin was reported [36]. Our data point the other way for WTAP, which we find elevated in SSc PBMCs alongside METTL3 and METTL14, and a two-fold rise in global m6A, suggesting that different tissue compartments, i.e. circulating immune cells versus whole lesional skin, a mixture dominated by fibroblasts, keratinocytes and infiltrating leukocytes, may respond differently to the disease. Our study substantially extends these observations by providing the first transcriptome-wide maps of m6A sites in SSc. Our findings are however consistent with reports from other fibrotic conditions. Increased expression of m6A writer components has been documented in lung samples from patients with idiopathic pulmonary fibrosis [24], liver samples from patients with hepatic fibrosis [25], and kidney tissue from patients with chronic kidney disease [26]. This convergence across different fibrotic diseases suggests that epitranscriptomic dysregulation, particularly involving m6A, may represent a common pathogenic mechanism in chronic fibrotic disorders.

Previous studies have shown a multi-layered interaction between the m6A machinery and TGF-β signaling, suggesting a bidirectional regulatory relationship. On one hand, TGF-β treatment of lung fibroblasts has been shown to induce expression of m6A writers [37], indicating that profibrotic stimuli can directly modulate the epitranscriptomic machinery. On the other hand, m6A-responsive sites are present in the 5’ untranslated region of *TGFB1* and control its expression under proinflammatory conditions [28], supporting a feedback mechanism whereby m6A can amplify TGF-β signaling. This bidirectional interaction may create a feed-forward loop that perpetuates fibrotic responses once initiated. The mechanistic connection between TGF-β signaling and m6A extends beyond transcriptional regulation. Moreover, downstream mediators of TGF-β signaling, such as SMAD2/3, physically interact with m6A writer enzymes guiding them to TGF-β-responsive transcripts, thereby affecting their stability or decay [38]. This direct recruitment mechanism provides spatial and temporal specificity to m6A deposition, ensuring that profibrotic transcripts are preferentially modified in response to TGF-β stimulation. Our observation that several genes harbour differentially methylated sites in SSc PBMCs is consistent with this targeted recruitment model. METTL3-mediated m6A deposition has been shown control the stability of *JUNB* and the translation efficiency of *JUN* [29,30], two transcription factors activated by TGF-β signaling that are central for skin and lung fibrogenesis [39,40]. In our Nanopore sequencing analysis, we identified methylation enrichment in the TGF-β signaling pathway, with JUNB as one of the hypermethylated genes in SSc PBMCs. Moreover, we detected differential methylation of KLF10, which was recently identified as a member of an inflammatory gene module along with JUNB in monocytes from SSc patients [34]. The methylation of KLF10 was exclusively detected in coding regions of SSc samples and correlated with increased transcript abundance, suggesting that m6A may enhance the expression of this profibrotic transcription factor through improved transcript stability or translational efficiency.

Here, we also reveal a distinct shift in m6A distribution patterns between SSc and healthy controls. While control samples displayed the expected enrichment of m6A around stop codons and 3′UTRs [23,24], regions classically involved in regulation of transcript stability, SSc samples showed increased m6A methylation in coding regions at the expense of 3′UTRs. This redistribution of m6A marks may reflect fundamental changes in how the m6A machinery recognizes and processes transcripts in the disease state. The increased prevalence of m6A in coding regions in SSc samples is particularly intriguing, as m6A modifications in these regions can affect translation elongation rates, ribosome pausing, and co-translational protein folding. This altered distribution pattern may represent an adaptive mechanism whereby cells fine-tune protein production in response to chronic stress and inflammatory signals. KLF10, where m6A methylation was detected exclusively in the coding region of SSc samples and correlated with increased transcript levels, may show how this altered topology may contribute to pathogenic gene expression programs.

Mesenchymal cell transition and activation of myofibroblasts are key pathogenetic processes in tissue fibrosis in SSc [5]. The conversion of tissue-resident fibroblasts into activated myofibroblasts is a hallmark of fibrotic tissue remodeling. Our findings demonstrate that pharmacological inhibition of METTL3 with STM2457 significantly attenuated TGF-β-induced expression of *ACTA2* and *COL1A1* in dermal fibroblasts, directly implicating m6A in the fibroblast-to-myofibroblast transition process. Recent studies have revealed tissue-specific mechanisms by which m6A promotes fibroblast activation. In the lung, METTL3-mediated m6A has been shown to promote fibroblast-to-myofibroblast transition by modulating translation of KCNH6 in a YTHDF1-dependent manner [27]. Similarly, m6A promotes mesenchymal transition of lung epithelial cells and lung fibrosis through modulation of JUN and JUNB [30]. Conversely, the m6A demethylase ALKBH5 inhibits TGF-β-induced mesenchymal transition of lung epithelial cells through demethylation and subsequent decreased transcript stability of TGFBR2 and SMAD3 [41], highlighting the importance of a balanced m6A homeostasis. Beyond the lung, m6A has been implicated in fibrogenesis across multiple organs relevant to SSc. METTL3 deficiency attenuates hepatic stellate cells activation and liver fibrosis [25], while targeting METTL3 has been shown to attenuate the development of kidney fibrosis through reduction of mesenchymal transition of renal epithelial cells and interstitial kidney fibrosis [26]. These observations suggest that m6A-mediated regulation of fibrotic processes may represent a conserved mechanism across different tissue types, which is particularly relevant for SSc given its multi-organ involvement [1]. Nonetheless, several considerations should guide future therapeutic development. First, while global inhibition of METTL3 effectively reduced profibrotic gene expression in our in vitro experiments without apparent toxicity, the safety profile of chronic m6A writer inhibition in vivo requires careful evaluation, given the fundamental role of m6A in normal cellular processes [9–11]. Second, the optimal timing of intervention, whether m6A-targeted therapy would be most effective in early inflammatory stages or later fibrotic stages, remains to be determined. Third, combination approaches pairing m6A inhibitors with existing immunosuppressive or antifibrotic therapies may prove more effective than monotherapy, particularly given the central role of TGF-β signaling in SSc pathogenesis [3,6,30]. Testing METTL3 inhibitors in preclinical models of dermal fibrosis is the necessary next step toward validating m6A as a therapeutic target.

Despite using two complementary platforms to detect m6A RNA methylation, our study has some limitations. Computational modification calling from direct RNA sequencing remains an evolving methodology in which false positives and platform-specific bias cannot be excluded, so site-resolved orthogonal validation, by m6A-sensitive reverse transcription or mass spectrometry, will be required for individual transcripts. Furthermore, our nanopore cohort comprised three individuals per group, adequate for discovery but not for effect-size estimation. Profiling was confined to PBMCs, chosen for accessibility and for the established contribution of circulating immune cells to SSc, but not the compartment in which fibrosis occurs since the epitranscriptomic landscape of lesional skin and lung may differ, and cell type–specific profiling of fibroblasts, endothelial cells and tissue-resident immune populations is needed to relate m6A signatures to local tissue damage. Finally, determining which readers transduce methylation into altered transcript fate at profibrotic genes and the driver beyond the increased writers expression would provide a more comprehensive understanding of how epitranscriptomic changes translate into altered cellular phenotypes and disease manifestations.

Overall, we provide the first comprehensive characterization of the m6A methylome in SSc, revealing widespread epitranscriptomic dysregulation characterized by increased expression of m6A writers, elevated global m6A levels, and altered distribution of m6A marks across the transcriptome. The enrichment of differentially methylated sites in TGF-β signaling pathway components, combined with the functional demonstration that METTL3 inhibition attenuates profibrotic gene expression, establishes m6A as a potentially important regulator of fibrotic responses in SSc. These findings not only expand our understanding of SSc pathogenesis but also identify the m6A modification machinery as a promising therapeutic target. As small-molecule METTL3 inhibitors advance through clinical development, our work provides a strong rationale for investigating these compounds in SSc and potentially other fibrotic diseases. The epitranscriptomic layer of gene regulation may represent an underexplored dimension of SSc pathobiology, and targeting this layer could offer novel approaches to interrupt the fibrotic cascade that drives disease progression.

## MATERIALS AND METHODS

### Patient recruitment

Thirty-one (31) SSc patients according to the 2013 ACR/EULAR classification criteria were recruited, while 24 apparently healthy individuals served as controls. Exclusion criteria included non-disease related severe renal or heart, cancer, recent infection (last 2 weeks). Demographics, clinical and laboratory features [interstitial lung disease, skin involvement, Raynaud’s phenomenon; puffy fingers; digital ulcers; erythrocyte sedimentation rate (ESR; mm/1^st^ h.)] and disease-specific auto-antibody status were recorded at baseline and have previously described. All participants gave informed consent in compliance with the Declaration of Helsinki, which had been previously approved by the Ethics Committee of Laiko Hospital, Athens, Greece (Protocol Nr.:1368/ 17-11-2016).

### Peripheral blood mononuclear cell isolation

Peripheral blood was collected in EDTA-containing tubes (BD Vacutainer). Peripheral blood mononuclear cells (PBMCs) were isolated within 2 hours by standard gradient centrifugation methods using Ficoll (Ficoll Paque Plus, Sigma Aldrich/Merck). Isolated PBMCs were lysed in Trizol and stored in a −80 freezer until further processing.

### Cell culture

A normal human dermal fibroblast cell line (HDFa; C0135C, Gibco) was cultured in Dulbecco’s Modified Eagle Medium (DMEM; 10569010, Gibco), supplemented with 10% Fetal Bovine Serum (FBS; Α5256701, Invitrogen) and 1% penicillin/streptomycin (P.S.; 15140122, Gibco). Cells were maintained at 37 °C in a humidified incubator with 5% CO₂.

Primary human dermal fibroblasts were isolated from forearm punch skin biopsies of patients with systemic sclerosis (SSc). Biopsy specimens were placed in tissue culture dishes and cultured in DMEM supplemented with 10% FBS, 3% fibroblast growth supplement (FGS; 116-GS, Sigma-Aldrich), 1% penicillin/streptomycin/amphotericin B (P/S/A; 15240062, Gibco), 0.1% gentamicin (10 mg/mL, 15710064, Gibco), and 1× non-essential amino acids (NEAA; 11140050, Gibco). After tissue adherence and fibroblast outgrowth, cells were transferred to tissue culture flasks and maintained in DMEM supplemented with 10% FBS and 1% penicillin–streptomycin. All cultures were maintained at 37 °C in a humidified atmosphere containing 5% CO₂.

### In vitro STM2457 treatment

HDFa cells and SSc-derived dermal fibroblasts were seeded at subconfluent level in DMEM, supplemented with 10% FBS and 1% penicillin/streptomycin. Twenty-four hours before treatment, cells were serum-starved in DMEM containing 1% FBS and 1% penicillin–streptomycin. Cells were treated with STM2457 (20um, HY-134836, MedChem Express) in the presence of recombinant human TGF-β1 (10 ng/mL; Human TGF-β1 Recombinant Protein, 100-21-10G, PeproTech) for 48 h. DMSO was used as a vehicle control and equal volumes were added to all experimental groups. Final DMSO concentrations did not exceed 0.5%, previously established to be non-toxic to the cells. All treatments were performed in starvation media (DMEM, 1% FBS, 1% P.S.) unless otherwise noted.

### Cell viability assay

Cell viability was assessed by crystal violet staining. Briefly, after the 48h treatment of the cells with STM2457, culture medium was removed and cells were washed once with phosphate-buffered saline (PBS) to remove non-adherent (dead/detached) cells. Cells were then fixed with ice-cold methanol for 15 min at room temperature and subsequently stained with 0.5% crystal violet solution for 10 min at room temperature under gentle agitation. Excess stain was removed by washing the plates four times with distilled water, and plates were air-dried overnight at room temperature. Bound crystal violet was then solubilized with 33% Acetic Acid for 5 min at room temperature under gentle agitation, and absorbance was measured at 570nm using a microplate reader. Absorbance values were normalized to the DMSO vehicle control, set as 100%, to determine relative cell viability across treatment conditions. All experiments were performed in triplicate.

### RNA extraction, reverse transcription, quantitative polymerase chain reaction

Total RNA was isolated from PBMCs using the Direct-zol RNA Miniprep Kit (R2052/R2053, Zymo Research) according to manufacturer’s instructions. One microgram of total RNA was reverse transcribed into complementary DNA (cDNA) using M-MLV reverse transcriptase (Invitrogen, ThermoFisher Scientific) and cDNA was diluted to a final volume of 200μl. Expression of *RPLP0* (housekeeping gene*; Fw: TCG ACA ATG GCA GCA TCT AC; Rv: ATC CGT CTC CAC AGA CAA GG*), *METTL3 (Fw: CAA GCT GCA CTT CAG ACG AA; Rv: GCT TGG CGT GTG GTC TTT)*, *METTL14 (Fw: AGA AAC TTG CAG GGC TTC CT; Rv: TCT TCT TCA TAT GGC AAA TTT TCT T),* and *WTAP (Fw: TTC CCA AGA AGG TTC GAT TG; TGC AGA CTC CTG CTG TTG TT)* was quantified with SYBR Premix Ex Taq (Takara) / PowerUP SYBR Green Master Mix (Applied Biosystems) on ViiA7 / QuantStudio 7 Flex system, respectively. The relative expression of each gene was determined according to the formula 2-ΔCt, where ΔCt = Ct (gene)-Ct (housekeeping gene).

Total RNA was isolated from fibroblasts using the Direct-zol RNA Miniprep Kit (R2062, Zymo Research) according to manufacturer’s instructions. One microgram of total RNA was reverse transcribed into complementary DNA (cDNA) using the PrimeScript™ RT Reagent Kit (Takara, Shiga, Japan) and cDNA was diluted to a final volume of 200μl. Expression of *RPLP0* (housekeeping gene*; Fw: TCG ACA ATG GCA GCA TCT AC; Rv: ATC CGT CTC CAC AGA CAA GG*), *ACTA2* (*Fw: CTA TGC CTC TGG ACG CAC AAC T; Rv: CAG ATC CAG ACG CAT GAT GGC A), KLF10 (Fw: AGG AGT CAC ATC TGT AGC CAC C; Rv: GAA CGG GCA AAC CTC CTT TCA C), COL1A1 (Fw: GAT TCC CTG GAC CTA AAG GTG C; Rv: AGC CTC TCC ATC TTT GCC AGC A)* was quantified with PowerUP SYBR Green Master Mix (Applied Biosystems) on QuantStudio™ 5 Real-Time PCR System, Applied Biosystems. The relative expression of each gene was determined according to the formula 2-ΔCt, where ΔCt = Ct (gene)-Ct (housekeeping gene).

### Global m6A quantification

Total N6-methyladenosine (m6A) RNA levels were quantified using the EpiQuik ™ m6A RNA Methylation Quantification Kit (P-9005, EpiGenTek) according to manufacturer’s instructions. 300ng total RNA were used as input per sample and were quantified according a standard curve generated by input samples with known m6A methylation concentration. All samples were run in the same plate to avoid inter-assay variability.

### Arraystar m6A Single Nucleotide Microarray

Total RNA from each sample was quantified using the NanoDrop ND-1000. The sample preparation and microarray hybridization were performed based on Arraystar’s standard protocols. Briefly, the total RNA was divided into two fractions. One fraction denoted as “Digested” was treated with RNA endoribonuclease MazF to cleave the unmodified m6A sites; the other fraction denoted as “Input” was untreated with MazF for both modified and unmodified sites. The “digested” RNAs were immunoprecipitated with anti-m6A antibody as “Modified” RNAs to enhance the m6A modified site signals. The “Modified” and “Input” RNAs were separately labeled with Cy5 and Cy3 as cRNAs using Arraystar Super RNA Labeling Kit. The “Modified” and “Input” cRNAs were combined together and hybridized onto Arraystar Human m6A Single Nucleotide resolution Microarray (8×15K, Arraystar). After washing the slides, the arrays were scanned in two-color channels by an Agilent Scanner G2505C.

Agilent Feature Extraction software (version 11.0.1.1) was used to analyze acquired array images. Raw intensities of Modified (Cy5-label) and Input (Cy3-label) RNAs were normalized with the average of log2-scaled Spike-in RNA intensities. After Spike-in normalization, the probe signals having Present (P) or Marginal (M) QC flags in at least 3 out of 13 samples were retained. “m6A site methylation stoichiometry” was calculated for the percentage of modification for each site and “m6A site abundance” was calculated for the m6A site methylation amount. Differentially m6A-methylated sites between two comparison groups were compiled by the fold change (>1.5) and statistical significance (p-value<0.05) thresholds. Hierarchical Clustering was performed to display m6A-methylation pattern among samples.

### Sample preparation and direct RNA sequencing

RNA from peripheral blood mononuclear cells (PBMCs) and control samples was processed for direct RNA sequencing. Libraries were prepared using the Oxford Nanopore SQK-RNA002 Direct RNA Sequencing Kit and sequenced on a PromethION platform using R9.4.1 flow cells (FLO-PRO002). Samples were subjected to quality control on the Agilent Bioanalyzer RNA Pico assay. As small RNAs are preferentially ligated during library preparation and can exhaust sequencing pores, an additional 2.2× bead-based clean-up step was introduced prior to library preparation to improve run performance and basecalling quality.

### Sequencing and base-calling

Real-time data acquisition and base-calling were performed with MinKNOW v22.12.5, Bream v7.4.8, MinKNOW Core v5.4.3, and Guppy v6.4.6 in high-accuracy mode, with a minimum quality score filter of Q ≥ 7. Sequencing runs generated both FAST5 (raw signal) and FASTQ (base-called reads) files. Read length distributions and q-score performance were monitored across runs, and sequencing yield exceeded 1 Gb for the majority of samples.

### Guided transcriptome assembly

All high-quality FASTQ reads were concatenated and aligned to the human reference genome (hg38, Ensembl GRCh38 release 104) using minimap2 (v2.24) with splice-aware alignment parameters (”-ax splice”). De novo transcriptome assembly was constructed by StringTie (v2.2.1) with the long reads processing mode (”-L”) and guided by the Ensembl GRCh38.104 annotation GTF, generating a custom GTF file. De novo cDNA sequences were extracted from the reference hg38 genome using “bedtools getfasta” (v2.30.0).

### Signal-level alignment and m6A site detection

For each sample, the base-called FASTQ reads were first aligned to the de novo transcriptome assembly using minimap2, then processed together with FASTQ reads, raw FAST5 signal data, and the reference de novo transcriptome uing “nanopolish eventalign” (v0.14) to resquiggle the raw signals and align individual signal events to the reference. To detect m6A modifications, program m6Anet (v2.0.0) [42] was used to process the output of nanopolish, and 1000 iterations were performed through each potential m6a sites during m6anet inference. We defined high confidence m6A sites as those with a probability modified value ≥ 0.9, for which the corresponding transcript was expressed and present in > 50% of samples expressed a probability that the site is modified in at least one group. To compare m6A sites between groups, we defined differential sites as those with a difference in modification ratio > 0.01 or < −0.01.

### RNA-sequencing of fibroblast transcriptome and data analysis

For transcriptome profiling, total RNA was isolated and used to prepare RNA-seq libraries using a DNBSEQ-compatible library preparation workflow according to the manufacturer’s instructions. Briefly, RNA was fragmented and reverse-transcribed into cDNA, followed by adapter ligation and library amplification. The resulting libraries were sequenced on a DNBSEQ-G400 platform. Raw sequencing reads were subjected to quality control by BGI, and gene-level raw read count matrices delivered by BGI were used as input for downstream analysis.

Differential gene expression analysis was performed using an edgeR/limma workflow as described previously [43]. Briefly, raw read counts were imported into R/Bioconductor and converted to counts per million (CPM) and log-CPM values using the cpm function from the edgeR package. Lowly expressed genes were filtered using the filterByExpr function in edgeR, excluding genes with fewer than 10 read counts in the required minimum number of samples. Batch effects were adjusted using the removeBatchEffect function applied to the log-CPM matrices. Heteroscedasticity inherent to RNA-seq count data was handled using the voom function from the limma package, which estimates the mean-variance relationship and calculates precision weights for each observation. Differential expression between control and treated samples was then assessed using limma linear modelling with empirical Bayes moderation. Log2 fold changes and Benjamini-Hochberg-adjusted p-values were calculated for each gene. Genes with an adjusted p-value <0.05 were considered significantly deregulated.

### Ingenuity Pathway Analysis

Ingenuity Pathway Analysis (IPA; QIAGEN, Hilden, Germany) was used to identify biological pathways, upstream regulators, molecular networks, and functional categories associated with the observed transcriptional changes. For each comparison, differentially expressed genes were uploaded into IPA together with their corresponding gene identifiers, log2 fold changes, and statistical significance values. Genes passing a p-value cutoff of <0.05 were included in the analysis. IPA Core Analysis was then performed using the default settings to evaluate enrichment of canonical pathways, biological functions, disease annotations, and predicted upstream regulatory mechanisms. The statistical significance of pathway enrichment was calculated by IPA using a right-tailed Fisher’s exact test, while activation z-scores, where available, were used to predict the direction of pathway or regulator activation.

### Statistical analysis

Statistical analysis was conducted with SPSS v26.0 and GraphPad Prism v10. Normality of continuous variables was assessed by D’Agostino-Pearson, Kolmogorov-Smirnov and Shapiro-Wilk tests. ROUT test (Q=1%) was used to identify outliers. Pairwise differences were evaluated with independent samples Student’s t-test (with Welch’s correction when variances of the groups were unequal) or with the non-parametric Mann-Whitney U test for continuous variables, and chi-squared test for nominal variables. Correlations between continuous variables were explored by Pearson’s correlation coefficient test. Results were considered statistically significant when P<0.05.

## Acknowledgments

The authors would like to thank Dr Kleio-Maria Verrou for help with initial bioinformatic analysis of microarray data.

## Funding

National Hellenic Research Foundation grant 28406 (NIV)

British Heart Foundation grant PG/23/11093 (ST-C)

Royal Society grant RG\R1\241197 (ST-C)

Program for Promoting Gender Equality and the Careers of Female Physicians and Scientists at University Medical Centre Mannheim, Medical Faculty Mannheim, Heidelberg University (KSo)

Special account for research grants, National and Kapodistrian University of Athens, Greece; grant 0974. (PPS)

German Research Foundation Deutsche Forschungsgemeinschaft, CRC1366 C07, project no. 394046768 (KSte)

Helmholtz Institute for Translational AngioCardioScience (HI-TAC) of the Max Delbrück Center for Molecular Medicine in the Helmholtz Association (MDC), Heidelberg University, Mannheim, Germany (KSte)

## Data and materials availability

All data needed to evaluate the conclusions in the paper are present in the paper and/or the Supplementary Materials. RNA sequencing data are deposited in the European Nucleotide Archive under accession number PRJEB113184. In accordance with donor consent and ethical approvals, the nanopore raw files can be provided pending scientific review and a completed data access agreement.

## Supplementary material

**Supplementary Figure 1.**
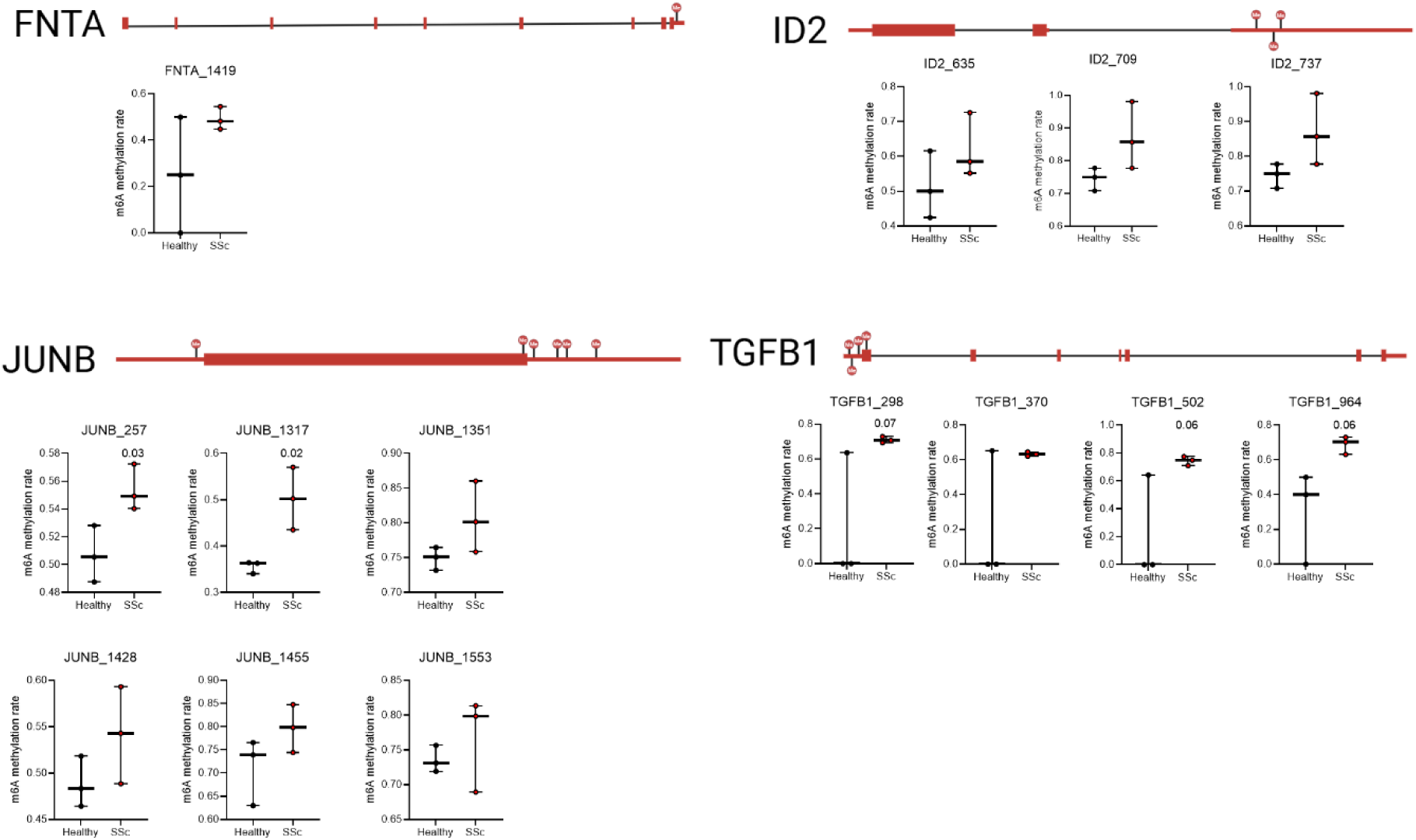
m6A modification along the transcript of TGF-b-related genes FNTA, ID2, JUNB and TGFB1 in HC and SSc PBMCs.

